# Gait transition mechanism from quadrupedal to bipedal locomotion in the Japanese macaque based on inverted pendulum

**DOI:** 10.64898/2025.12.07.692854

**Authors:** Koki Nishizaki, Mau Adachi, Yuichi Ambe, Yushi Tsuruse, Ryo Iba, Hiroko Oshima, Takashi Suzuki, Yasuo Higurashi, Kei Mochizuki, Katsumi Nakajima, Shinya Aoi

## Abstract

The ability of non-human primates to transition from quadrupedal to bipedal locomotion offers critical insights into both the evolution of human bipedalism and the principles of complex motor control. While quadrupedal and bipedal gaits in non-human primates have been studied, the dynamic mechanisms underlying the transition between these gaits remain poorly understood. Japanese macaques trained to walk bipedally have been reported to utilize inverted pendulum dynamics to achieve efficient bipedal locomotion. Given the intrinsic instability of inverted pendulum systems, which can induce large changes in movement with minimal control input, we hypothesized that this mechanism also contributes to the gait transition. To test this, we developed a neuromusculoskeletal model of the Japanese macaque that integrates a detailed musculoskeletal structure with a physiologically inspired motor control system. Through forward dynamics simulations, we generated a variety of movement patterns by systematically parameterizing motor commands, including failed transitions that are difficult to capture experimentally. We then applied dynamical systems analysis using an inverted pendulum model to examine the underlying principles of the transition process. Our results demonstrate that successful gait transitions depend on generating an inverted pendulum motion through appropriate control of the forward step length of one hindlimb. These findings provide mechanistic insights into how Japanese macaques coordinate their complex musculoskeletal systems to perform skilled, full-body movements in the gait transition, offering a deeper understanding of both advanced motor control and the evolution of human bipedalism.

## 1 Introduction

Humans have evolved to walk bipedally, but the origins of this are not well understood. On the other hand, non-human primates that walk not only quadrupedally but also bipedally provide important insights into the evolution of human bipedalism [1, 22, 40]. In the transition from quadrupedal to bipedal locomotion, some cases involve stopping and standing up before beginning to walk bipedally, while in others, the trunk is raised during continued movement, leading directly into bipedal walking. The former is a relatively static transition, whereas the latter requires a dynamic shift from the relatively stable motion of quadrupedal walking to the inherently unstable nature of bipedal locomotion without losing balance. This dynamic transition is a complex, full-body movement that requires precise coordination of the limbs and trunk, as well as effective balance control. The acquisition of bipedalism involves not only the relatively static transition ability of the former, but also the highly dynamic transition ability of the latter. Understanding the mechanism of highly dynamic transition will lead to an understanding not only of the evolution of human bipedalism, but also of the superior motor control of animals. Although quadrupedal and bipedal locomotion of non-human primates has been well studied [8, 11, 14, 17, 26, 28, 35, 44], little research has been done on dynamic gait transitions, and the mechanism remains largely unclear.

Japanese macaques (*Macaca fuscata*) that have been trained to walk bipedally show some of the characteristics of human bipedal walking. In particular, they use an inverted pendulum mechanism [5], which explains the energy saving mechanism during walking by exchanging potential and kinetic energy based on inverted pendulum dynamics [16, 29, 31]. While a normal pendulum has a stable equilibrium at its downward position, an inverted pendulum has an unstable equilibrium at its upright position. The dynamics in the vicinity of this upright equilibrium are governed by saddle-type instability. This property enables the system to generate dynamic motion and induce large changes in movement with minimal control input. In the transition from quadrupedal to bipedal locomotion, the forelimbs are released from supporting the body and the body is supported only by the hindlimbs, like an inverted pendulum. The gait transition is completed by raising the trunk while walking on the hindlimbs. Therefore, we hypothesize that trained Japanese macaques exploit inverted pendulum dynamics to perform the dynamic gait transition from quadrupedal to bipedal locomotion.

The typical approach to understanding motor control in animals involves measuring their movements and analyzing the resulting data. However, unless special techniques or equipment are employed, this approach can generally only capture relatively standard or successful movements, making it difficult to investigate what occurs when animals deviate from these patterns. In particular, it is difficult to induce situations in Japanese macaques where gait transitions fail, such as through falling, which limits our ability to understand the underlying mechanisms of their dynamic gait transition. To overcome these limitations, forward dynamics simulations using neuromusculoskeletal models, which integrate detailed musculoskeletal models with motor control models based on neurophysiological evidence, have gained attention. This constructivist approach has been applied to uncover the mechanisms of animal locomotion [9, 12, 19, 43, 45, 46, 49], and has also been used to study the evolution of human bipedalism from an anthropological perspective [27, 32, 34, 41, 42].

In the present study, we build on this framework by developing a neuromusculoskeletal model of the Japanese macaque that incorporates a physiologically inspired motor control system governing all four limbs and trunk. By parameterizing this motor control model, we generated a variety of movement patterns, including successful transitions and failures such as falling. Using forward dynamics simulation and dynamical systems analysis, we tested the hypothesis and examined how Japanese macaques leverage inverted pendulum dynamics to achieve successful gait transitions. Our approach provides mechanistic insights into how Japanese macaques control their complex musculoskeletal systems to perform highly skilled, full-body movements, offering a deeper understanding of both skillful motor control and the evolution of human bipedalism.

## Results

### Neuromusculoskeletal model

We developed a two-dimensional musculoskeletal model of Japanese monkeys composed of eleven rigid links (one for the trunk, two for each forelimb, and three for each hindlimb) and 28 principal muscles (six for each forelimb and eight for each hindlimb) as shown in Fig. 1, which are driven by command signals from the motor control model.

**Figure 1:**
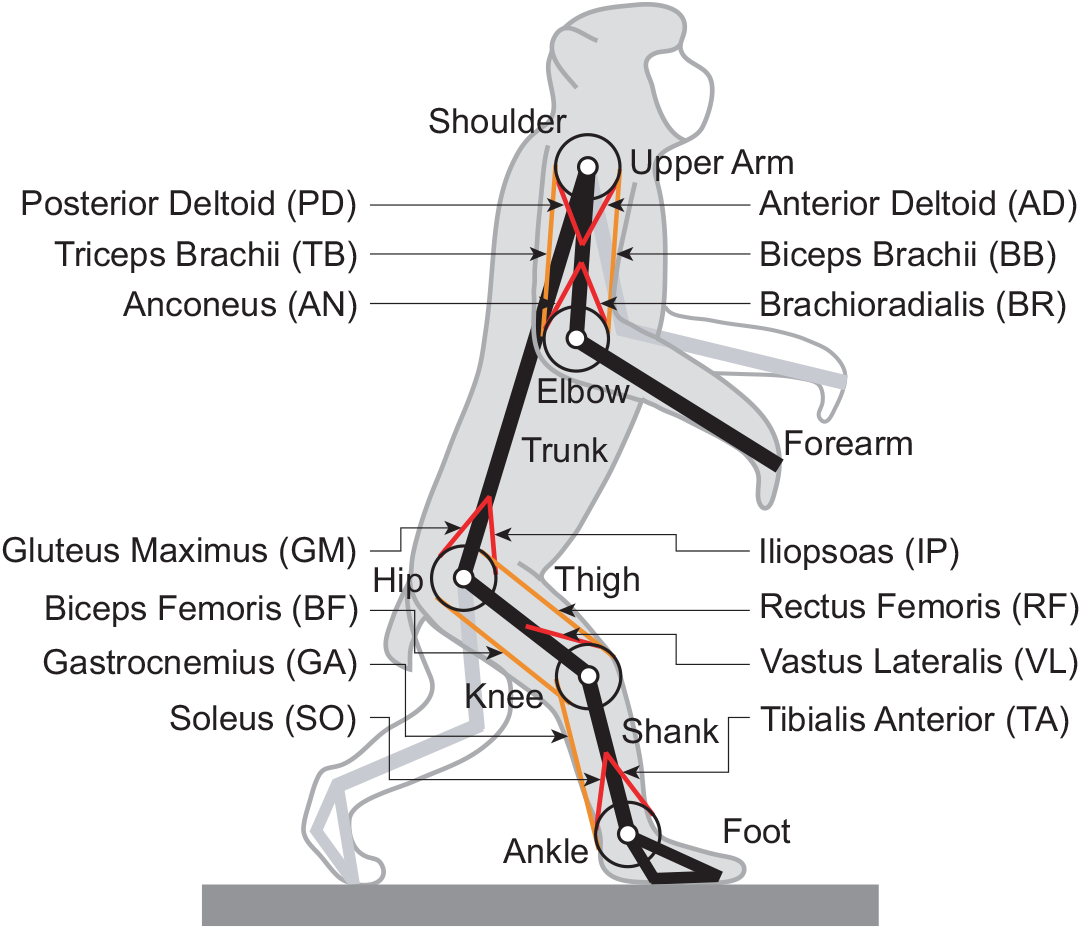
Musculoskeletal model of Japanese monkey.

The motor control model consists of two components. The first is the movement generator, which produces a small number of weighted activation pulses (rectangular pulses) in a feedforward manner for quadrupedal and bipedal walking and gait transition using phase oscillators based on the two-layer central pattern generator (CPG) model [4, 38] and the muscle synergy hypothesis [7,10,13,20,47]. The second is the movement regulator, which modulates locomotor activity in a feedback manner based on sensory information and reference values. The motor commands generated by the motor control model are transformed into muscle activities through a low-pass filter.

The movement generator for quadrupedal and bipedal walking uses four rectangular pulses corresponding to early extension, late extension, early flexion, and late flexion to generate periodic limb movements (Fig. 2). Although the same four pulses are used for both quadrupedal and bipedal walking, their onset and offset phases, as well as their weightings, differ between the two gaits. The gait cycle duration was set to 0.76 s for quadrupedal walking and 0.6 s for bipedal walking from the measured monkey data [15]. The movement regulator regulates the forward speed, hip height, and trunk posture to stabilize the locomotor behavior. Specifically, the forward speed is regulated by the IP, GM, TA, and SO muscles; the hip height by the VL, TA, and SO muscles; and the trunk posture by the IP and GM muscles (Fig. 3).

**Figure 2:**
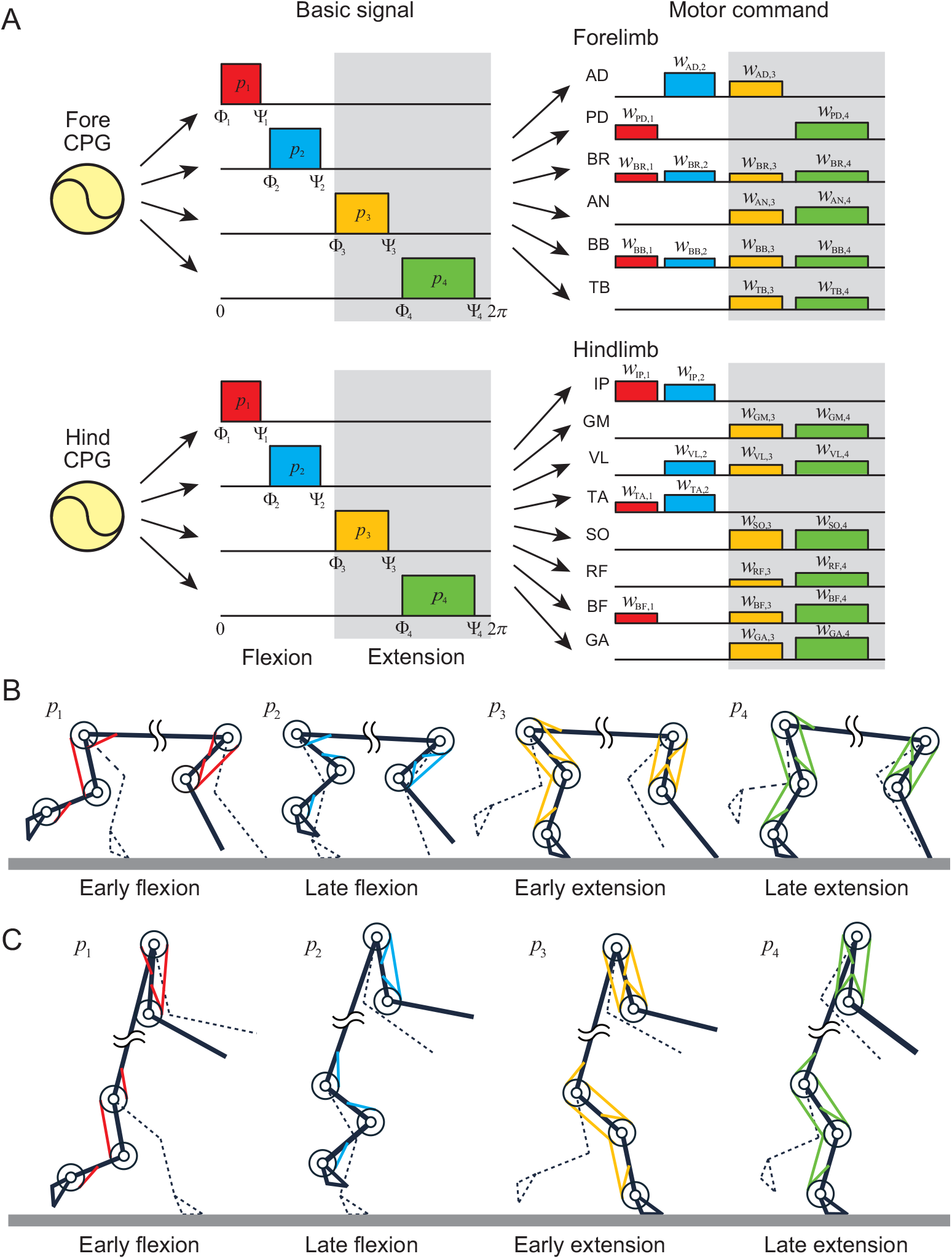
Movement generator of the motor control model based on the muscle synergy hypothesis. A. Motor commands represented as a linear combination of four basic signals. Although the same four signals are used for both quadrupedal and bipedal walking, their onset and off-set phases, as well as their weightings, differ between the two gaits. B. Coordinated muscles associated with each basic signal during quadrupedal walking. C. Coordinated muscles during bipedal walking. The colors used for the motor commands and muscles correspond to those of the associated basic signals. Note that the forelimb and hindlimb illustrations represent their respective phases and are not necessarily shown at the same instant.

**Figure 3:**
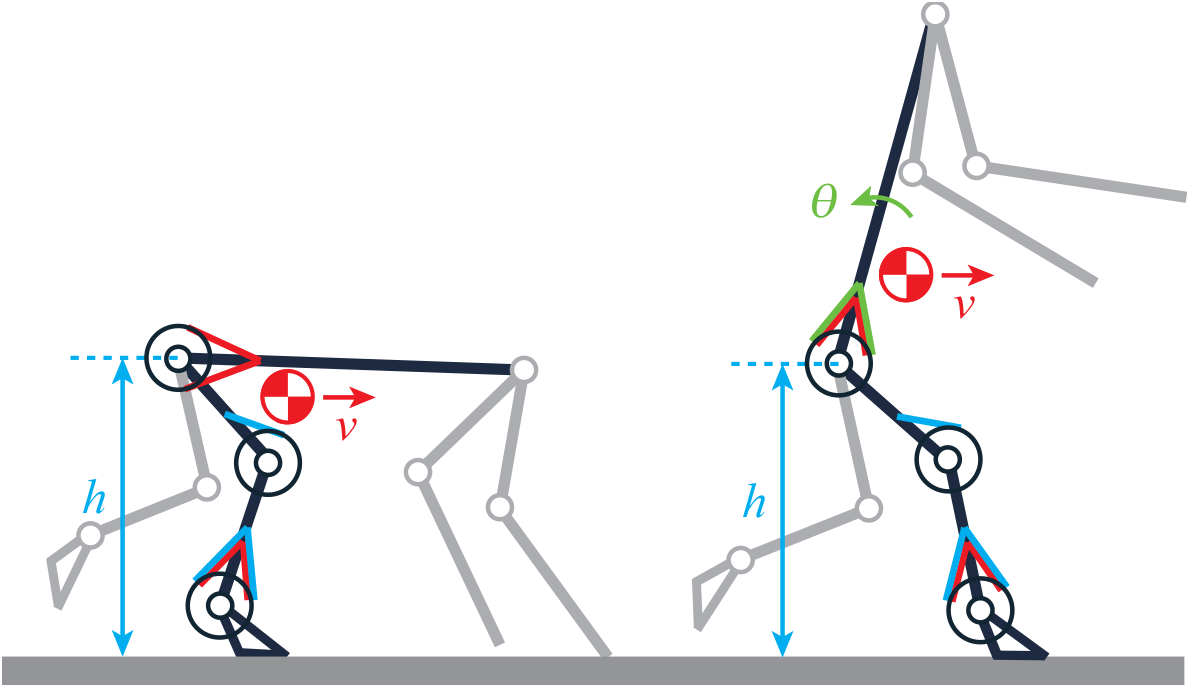
Movement regulator of the motor control model based on forward speed, hip height, and trunk posture (trunk posture is regulated only during bipedal walking). The control target and the muscles used for the control are shown in the same color.

For the gait transition from quadrupedal to bipedal walking, the movement generator adds two sequential rectangular pulses to the four rectangular pulses shown in Fig. 2 in the right hindlimb. One pulse is used to generate a large step in preparation for the gait transition, whereas the other is used to raise the trunk and execute the transition (Fig. 4). After the additional pulses are applied, the parameters of the motor control model are switched from those for quadrupedal walking to those for bipedal walking. Since the gait transition is achieved through the control of the right hindlimb, we refer to it as the trigger limb, and the other hindlimb as the trailing limb.

**Figure 4:**
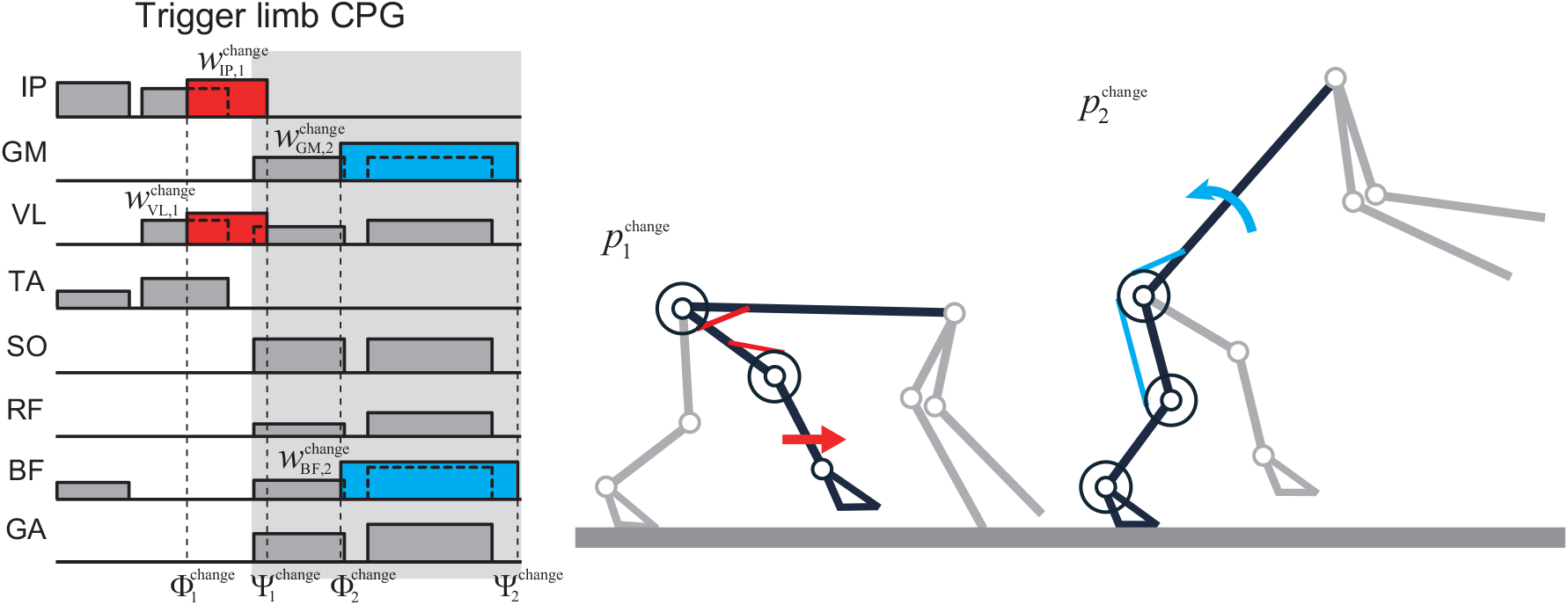
Gait transition by using two additional pulses to trigger limb muscles.

### Reproduction of quadrupedal and bipedal walking

The neuromusculoskeletal model successfully reproduced both quadrupedal and bipedal walking at speeds of 0.78 and 0.73 m/s, respectively (see Supplementary Movies S1 and S2). The parameters of the motor control model were determined using the particle swarm optimization (PSO) method [21], such that the error between the simulated and measured kinematic data was minimized, as summarized in Tables S1 and S2 in Supplementary Information S1.

Figure 5 compares the simulation results with the monkey data. Figures 5A and B show the stick diagram. Figures 5C and D show the elevation angle and muscle activity, respectively, for one gait cycle. The contribution of the movement regulator is much smaller than that of the movement generator, indicating that the movement generator accounts for most of the motor commands. During quadrupedal walking, the forelimbs and hindlimbs coordinate with each other to move forward. During bipedal walking, the trunk is maintained in an upright posture, and the forelimbs are released from body support. Although not all of the muscles incorporated in our model were measured in the monkey, and the simulation results show some discrepancies with the monkey data, such as a small thigh angle and a short double stance phase duration, the simulated results successfully reproduce the fundamental locomotor characteristics observed in the monkey. In particular, the trunk remains nearly horizontal during quadrupedal walking and upright during bipedal walking. The hindlimb elevation angles are coordinated such that they increase during the swing phase and decrease during the stance phase in both quadrupedal and bipedal walking. Furthermore, the trajectories exhibit similar patterns across the two gaits. The forelimb muscles exhibit little activity during bipedal walking. In contrast, the hindlimb muscles show greater activity, characterized by longer activation durations and larger activation amplitudes than those observed during quadrupedal walking.

**Figure 5:**
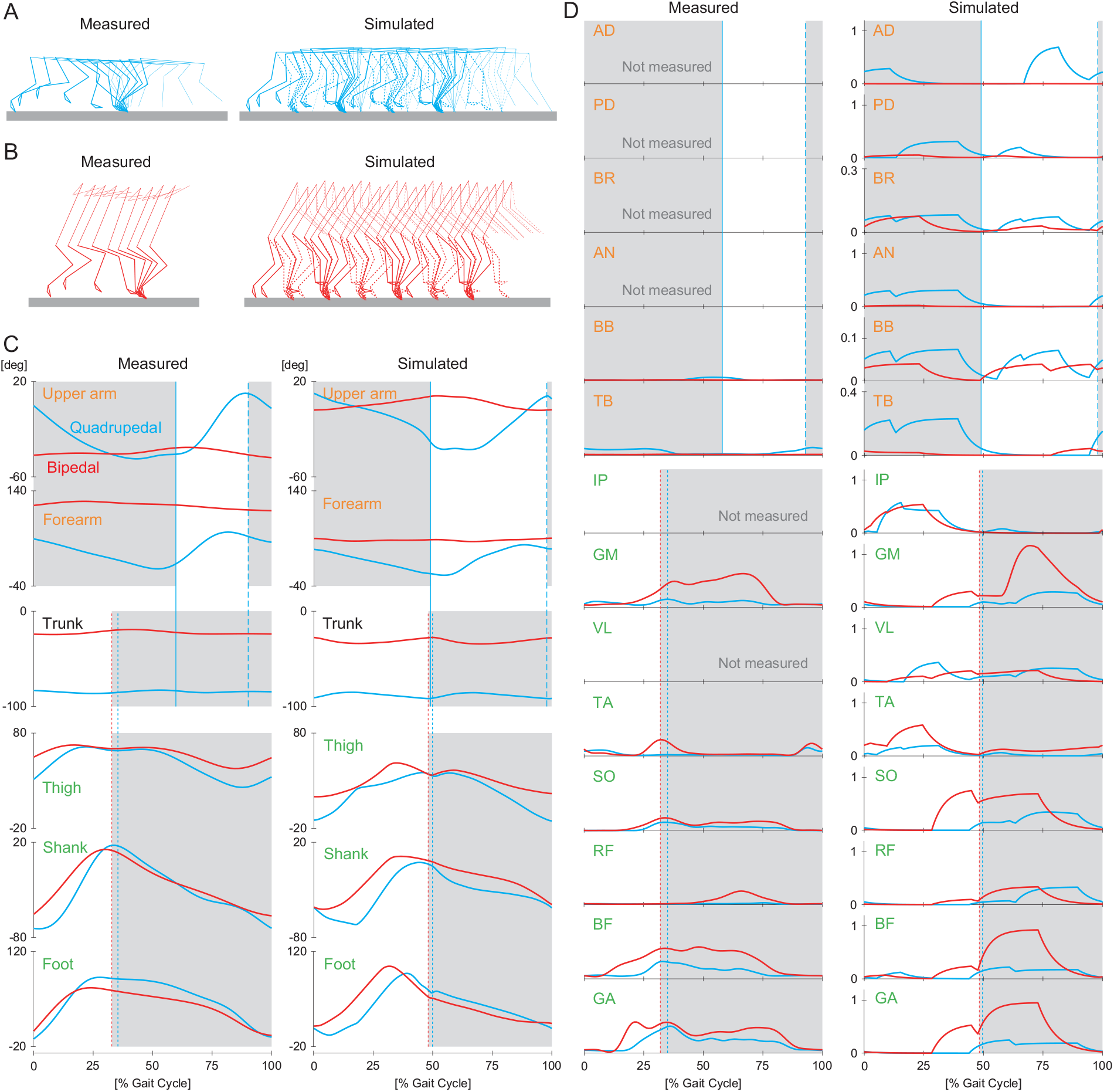
Simulation results of quadrupedal and bipedal walking compared with measured monkey data. A. Stick diagram of quadrupedal walking. B. Stick diagram of bipedal walking. In A and B, simulation results are shown for both sides (see Supplementary Movies S1 and S2), whereas measured data are shown only for the right side. C. Elevation angles. D. Muscle activity. In C and D, both simulation results and measured data are shown for the right side during quadrupedal (blue) and bipedal (red) walking. 0 and 100% of gait cycle indicate hindlimb liftoff. Vertical dotted, dashed, and solid lines indicate hindlimb touchdown and forelimb touchdown and liftoff, respectively. Gray regions indicate stance phase of each limb.

### Reproduction of gait transition

The model successfully changed the gait from quadrupedal to bipedal walking by using two additional signals in the trigger limb (see Supplementary Movie S3). The model parameters were determined using the PSO method to minimize the error between the simulated and measured kinematic data during bipedal walking after the gait transition, as shown in Table S3 in Supplementary Information S1. We also performed a sensitivity analysis of the optimized parameters and confirmed the robustness of the simulation results (see Supplementary Information S2 for details).

Figure 6 compares the simulation results with the monkey data. Figures 6A, B, and C show the stick diagram, elevation angle, and muscle activity, respectively. The monkey took a large step in the trigger limb by increasing the elevation angles of the thigh and shank, and then raised the trunk by increasing the activity of the GM and BF muscles. Although the simulation results show some discrepancies with the monkey data, such as the absence of large activity of the TA and SO muscles at touchdown during the transition, which would increase ankle joint stiffness through co-contraction, they still show similar patterns to those of the monkey during the gait transition.

**Figure 6:**
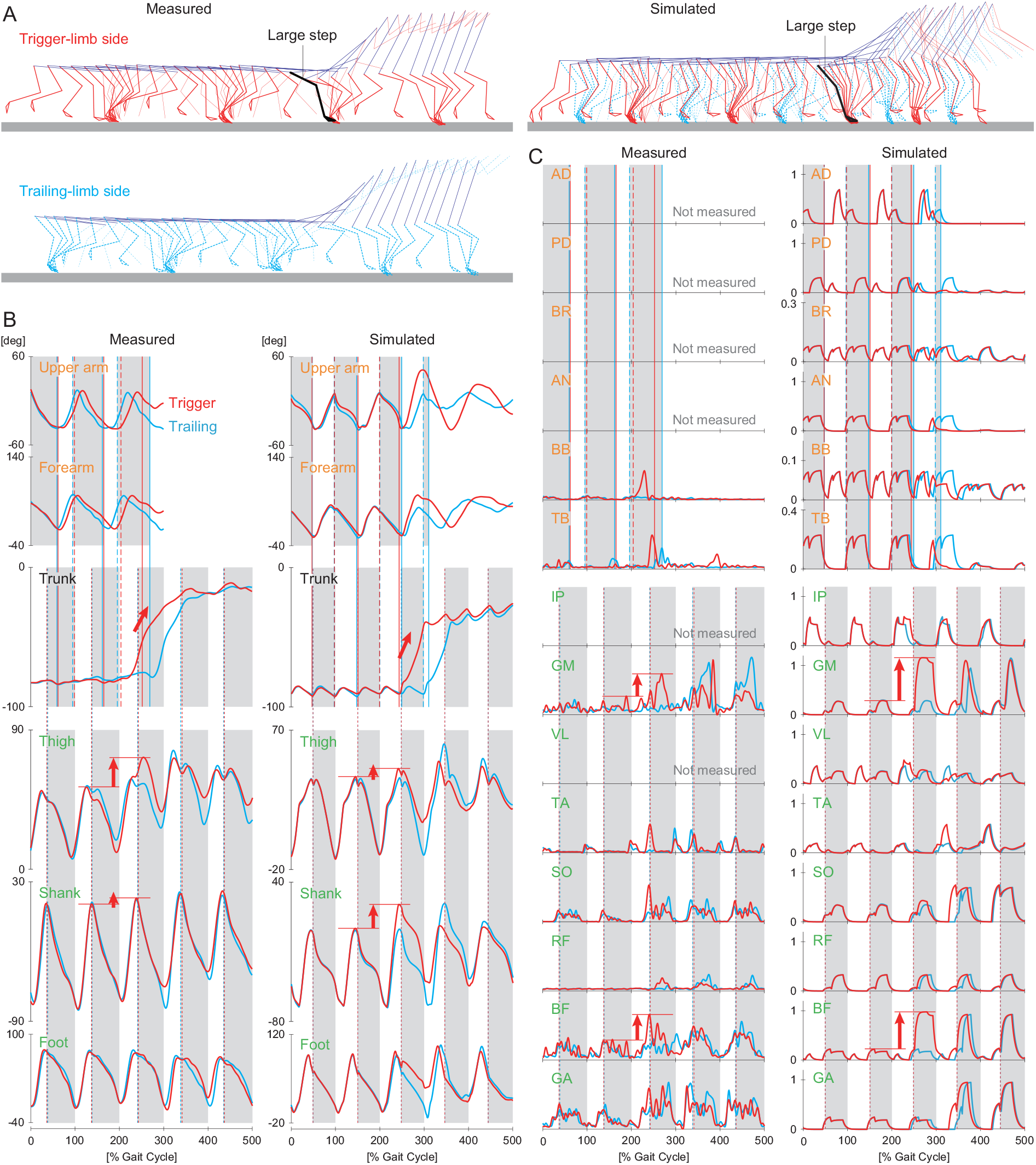
Simulation results of the transition from quadrupedal to bipedal walking compared with measured monkey data. A. Stick diagram (see Supplementary Movie S3). Measured data are shown separately for the trigger-limb and trailing-limb sides. B. Elevation angle. C. Muscle activity. These data are shown for the ipsilateral side of the trigger limb (red) and trailing limb (blue). 0, 100,, · · · 500% of gait cycle indicate hindlimb liftoff. Vertical dotted, dashed, and solid lines indicate hindlimb touchdown and forelimb touchdown and liftoff, respectively. Gray regions indicate stance phase of each limb. Elevation angles of the thigh and shank of the trigger limb are increased to take a large step before the transition, and then the trunk is raised by increasing the activity of the GM and BF muscles. These changes are indicated by red arrows.

### Contribution of the trigger limb to gait transition

To investigate the contribution of the large step generated by the trigger limb to the gait transition, we simulated the gait transition while varying the weighting coefficients 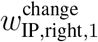 and 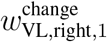 of the IP and VL muscles for the first additional pulse (Fig. 4). Specifically, these two weighting coefficients were systematically varied between 0 and 1, and the remaining parameters were reoptimized for each pair of coefficient values. We judged the simulated gait transition to be successful based on two criteria. The first criterion is if the center of mass (CoM) height averaged over one gait cycle after adding the two pulses is greater than 0.4 m. When normalized by the hindlimb length (0.42 m), this corresponds to a threshold of approximately 0.95 (≃ 0.4*/*0.42). The other criterion is if the trigger limb lifts off the ground within one gait cycle (0.6 s) after touchdown to continue walking. When normalized by the gait cycle duration, this corresponds to a threshold of 1.

Figure 7A shows the results for the horizontal distance between the CoM and the foot contact point normalized by the stride length (moving distance from one touchdown to the next) during quadrupedal walking before the transition, evaluated across the two parameters. This CoM-foot distance increases monotonically as the two parameters increase. Figure 7B shows the results for the first criterion, i.e., the CoM height averaged over one gait cycle after the two pulses were added, normalized by the hindlimb length. The area with a height greater than the success threshold is mainly divided into two regions on the left and right. Figure 7C shows the results for the second criterion, i.e., the time taken from touchdown to liftoff of the trigger limb when it took a large step, normalized by the gait cycle duration. The time suddenly increases on the right side and exceeds the success threshold. This indicates that although the CoM height is greater than the threshold on the right region of Fig. 7B, the transition failed (only the left region of Fig. 7B is successful).

**Figure 7:**
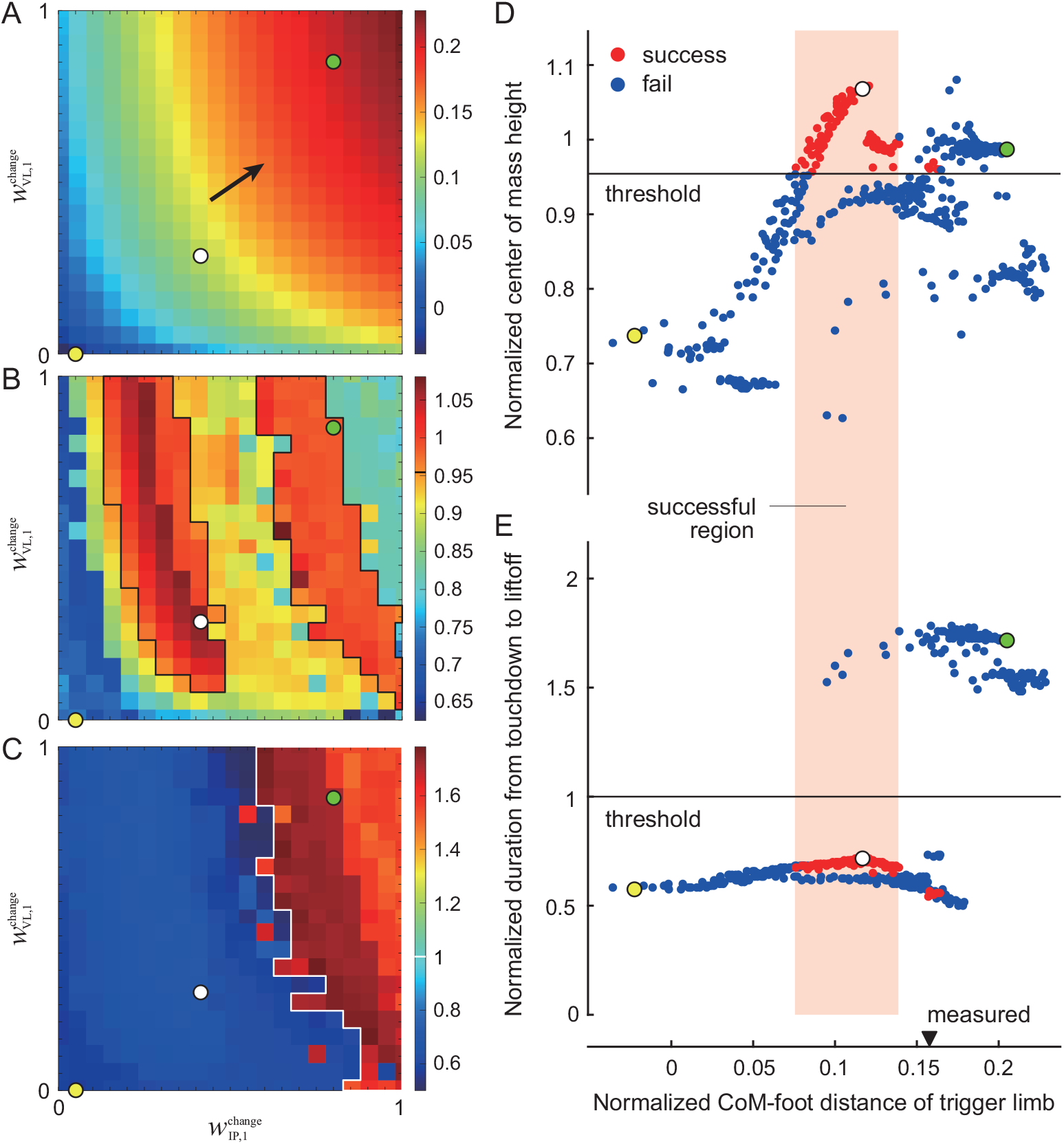
Investigation of contribution of trigger limb to gait transition using parameters 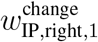 and 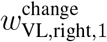. A. CoM-foot horizontal distance at touchdown of the trigger limb, normalized by the stride length during quadrupedal walking before the transition. B. First criterion: CoM height averaged over one gait cycle after adding two pulses, normalized by the hindlimb length. Black lines indicate the success threshold boundary (0.95 ≃ 0.4*/*0.42). C. Second criterion: Time from touchdown to liftoff of the trigger limb following a large step, normalized by the gait cycle duration. White lines indicate the success threshold boundary (1 = 0.6*/*0.6). From A, B, and C, the two criteria depending on the CoM-foot distance of the trigger limb are shown in D and E. White circles indicate the optimal result. Yellow and green circles indicate the examples used for comparison in Fig. 8. Successful gait transition is almost limited to a specific range (orange) of the CoM-foot distance of the trigger limb. The CoM-foot distance of the trigger limb is compared with the measured monkey data (arrow head).

Figures 7D and E reconstruct the results of Figs. 7A, B, and C. Specifically, they show two evaluation quantities in Figs. 7B and C (CoM height and time taken from touchdown to liftoff of the trigger limb) as a function of the CoM-foot horizontal distance at touchdown of the trigger limb in Fig. 7A. When the gait transition was successful, the CoM-foot distance was confined within a specific range, with only a few exceptions (see below for the reason). When the distance was greater or less, the transition failed. Figure 7E also shows the CoM-foot distance of the trigger limb calculated from the measured monkey data, indicating that the Japanese monkey took a slightly larger step during the transition than our model. This discrepancy is likely attributable to the fact that Japanese monkeys flex their trunk and employ pelvic yaw rotation when stepping forward.

### Hypothetical role of large trigger-limb step based on inverted pendulum model

When the trunk is raised during the gait transition, the forelimbs are free from supporting the body and bipedal walking begins with alternating left-right hindlimb stepping. Therefore, to clarify the successful gait transition mechanism depending on the forward step length of the trigger limb, we used an inverted pendulum model (Fig. 8A), whose angle is *θ*, connected from the touchdown position of the trigger limb to the CoM. This model follows the saddle dynamics and the motion changes greatly at the boundary between stable and unstable manifolds (Fig. 8B). Specifically, when the trajectory starts above the stable manifold in the left half plane, the model falls forward (Fig. 8B, a-c). In contrast, when the trajectory starts below the stable manifold, the model falls backward (Fig. 8B, d-f). Therefore, falling forward through the inverted pendulum motion with the support of the trigger limb and then landing the trailing limb in a suitable position to begin a new inverted pendulum motion would be a key to a successful transition.

**Figure 8:**
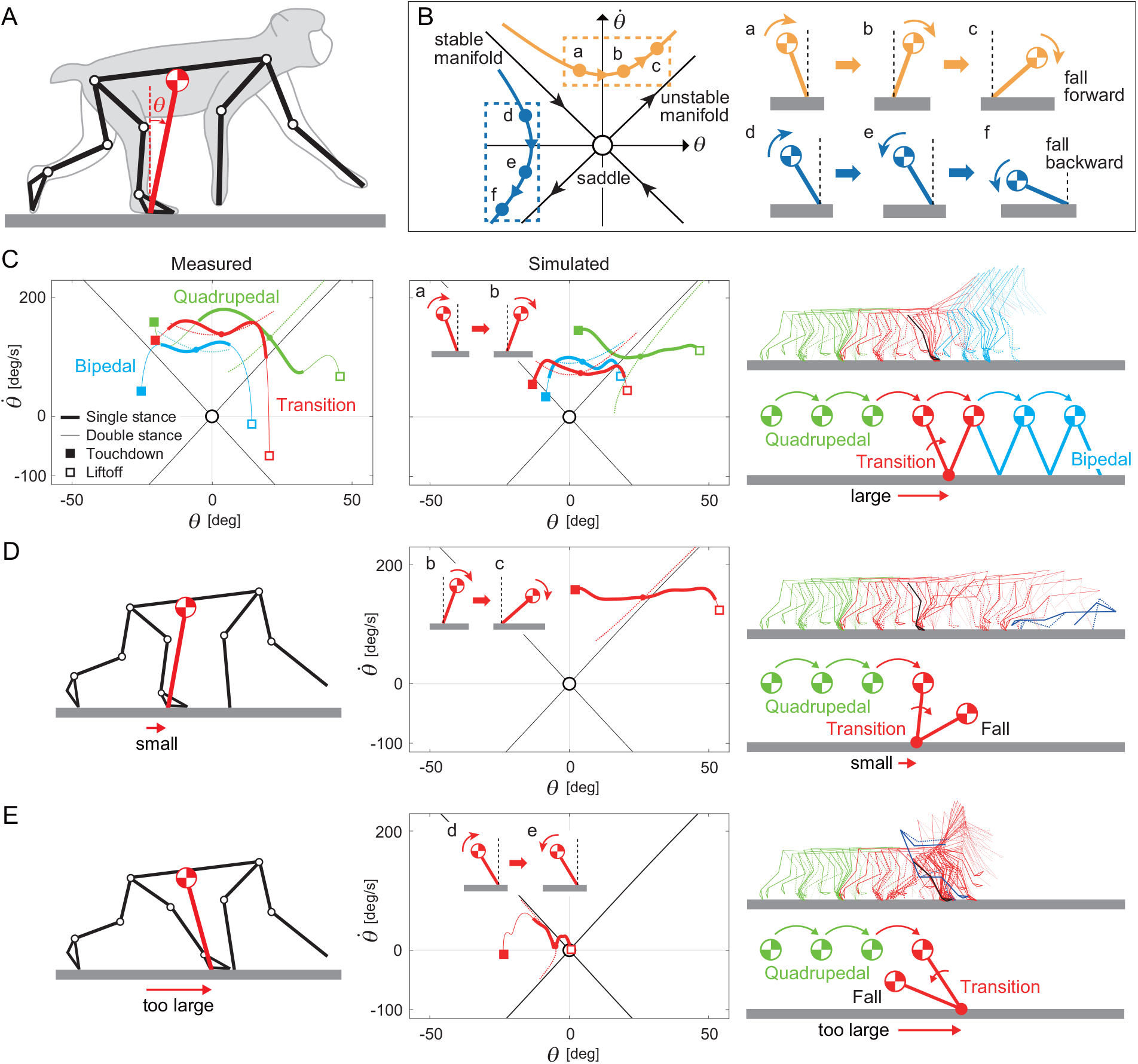
Role of forward step length of trigger limb in gait transition based on inverted pendulum model. A. Inverted pendulum model. B. Saddle dynamics. C. Application of simulation result of successful gait transition to inverted pendulum model, compared with application of measured monkey data. Dotted lines indicate the trajectories of rigorous inverted pendulum model. ‘Single stance’ and ‘double stance’ refer to the single and double stance phases of the hindlimbs, respectively. For both simulated and measured results of quadrupedal walking, the CoM position during the single stance phase starts just above the touchdown point of the trigger limb (*θ* ≃ 0) and moves forward. In contrast, during bipedal walking and gait transition, it starts behind the touchdown position (*θ <* 0) and advances forward through an inverted pendulum motion. D. Application of simulation result of failed gait transition due to small forward step length (see Supplementary Movie S4). E. Application of simulation result of failed gait transition due to too large forward step length (see Supplementary Movie S5).

First, Fig. 8C shows the result of applying the simulation result of the successful gait transition in Fig. 6 (indicated by the white circle in Fig. 7 with 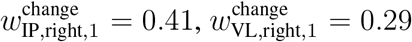) to the inverted pendulum model. This figure uses the simulation result from the touchdown to liftoff of the trigger limb after taking a large step. This figure compares the result during the gait transition with those during quadrupedal and bipedal walking in Fig. 5. This figure also compares the result of applying the simulation results to the inverted pendulum model with the result of applying the measured monkey data to the inverted pendulum model. We focused on the single stance phase of the hindlimbs to examine whether the CoM motion follows inverted pendulum dynamics. Although the trajectories during quadrupedal walking do not match those of an inverted pendulum due to the support by the forelimbs, they show that the CoM position starts just above the touchdown position of the trigger limb (*θ* ≃ 0) and moves forward. In contrast, the trajectories during bipedal walking are similar to those of an inverted pendulum, showing that the CoM position starts behind the touchdown position (*θ <* 0) and the model falls forward through the inverted pendulum motion. However, bipedal walking continues by switching to the inverted pendulum motion of the contralateral hindlimb. The trajectories during gait transition are similar to those during bipedal walking. This means that the trigger limb takes a larger step during the gait transition than it does during quadrupedal walking, which creates an inverted pendulum motion similar to that during bipedal walking and enables a successful gait transition. These characteristics during quadrupedal walking, bipedal walking, and gait transition are qualitatively similar between the simulation results and the measured monkey data.

Second, Fig. 8D shows the result of applying the simulation result of the failed gait transition due to small forward step length in Fig. 7 (indicated by the yellow circle in Fig. 7 with 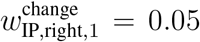 and 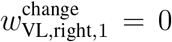, see Supplementary Movie S4) to the inverted pendulum model. The trajectory is similar to that during quadrupedal walking in Fig. 8C. In particular, the starting position of the trajectory is almost above the origin and much more to the right than that of the successful transition in Fig. 8C. This causes the model to fall forward rapidly, resulting in a failed gait transition.

Finally, Fig. 8E shows the result of applying the simulation result of the failed gait transition due to too large forward step length in Fig. 7 (indicated by the green circle in Fig. 7 with 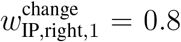 and 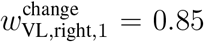, see Supplementary Movie S5) to the inverted pendulum model. The trajectory is below the stable manifold, which causes the model to fall backward. However, it should be noted that because the trailing limb remains in contact with the ground without lifting, the double stance phase continues for a while so that the model does not fall backward immediately and the CoM reaches a high position. This is the reason why, although the CoM height is greater than the threshold on the right region of Fig. 7B, the transition failed.

### Verification of hypothetical role of large trigger-limb step

Based on the results of the previous section, it is expected that the gait transition is successful through an inverted pendulum motion with appropriate control of the forward step length of the trigger limb. To verify this hypothesis, we performed detailed analyses based on the inverted pendulum model using the simulation results of the parameter study in Fig. 7. First, Fig. 9A shows the results of applying the simulation results during the gait transition to the inverted pendulum model, indicating that the trajectories move from the top right to the bottom left as the forward step length of the trigger limb increases.

**Figure 9:**
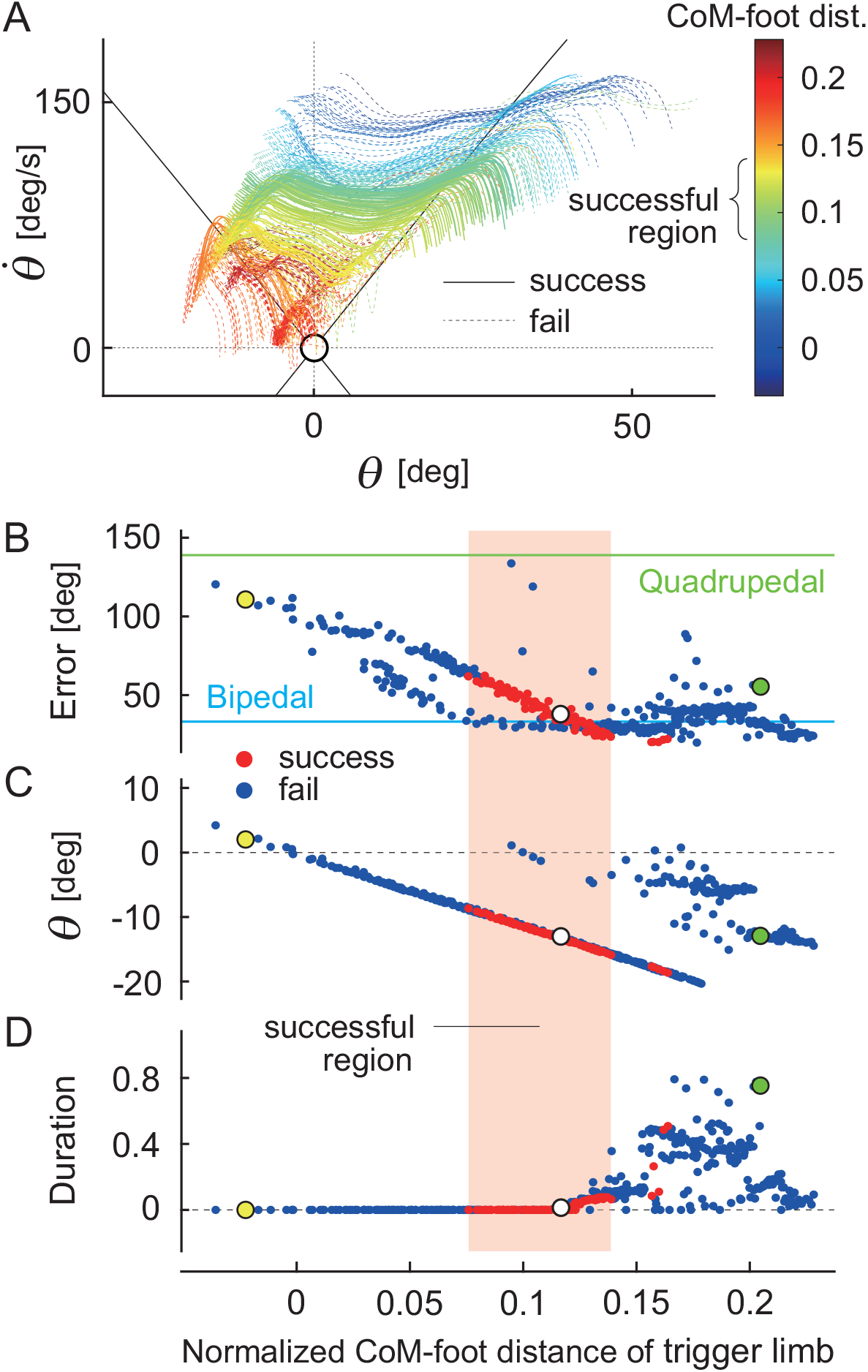
Verification of role of large forward step length of trigger limb in gait transition. A. Trajectories by applying simulation results for single stance phase of the hindlimbs during gait transition to inverted pendulum model, with the CoM-foot horizontal distance at touchdown of the trigger limb normalized by the stride length during quadrupedal walking before the gait transition. B. Root mean square error between the trajectories of simulation results and those of rigorous inverted pendulum model for t<u>he C</u>oM-foot distance of the trigger limb. The trajectories were nondimensionalized using 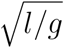, where *l* is the pendulum length and *g* is the acceleration of gravity. C. Onset angle *θ*. D. Duration when the trajectory is below the stable manifold, normalized by the gait cycle duration. White circles indicate the optimal result (Fig. 8C). Yellow and green circles indicate the results of Figs. 8D and E, respectively.

Second, we investigated whether the simulated monkey model exhibits inverted pendulum motion during the gait transition. Specifically, we examined the root mean square errors between the trajectories in Fig. 9A and those of the rigorous inverted pendulum model obtained by solving the equation of motion and compared them with the corresponding errors for quadrupedal and bipedal walking. Figure 9B shows the results for the CoM-foot horizontal distance at touchdown of the trigger limb, normalized by the stride length during quadrupedal walking before the gait transition. Although the error during the gait transition is larger than that for bipedal walking, it is much smaller than that for quadrupedal walking and tends to decrease as the forward step length of the trigger limb increases. This suggests that the monkey model exhibits more pronounced inverted pendulum motion during the gait transition as the forward step length increases.

Next, Fig. 9C shows the onset angle *θ* of the trajectories in Fig. 9A for the CoM-foot horizontal distance at touchdown of the trigger limb. This indicates that when the forward step length is small, the onset position of the CoM is above or to the right of the touchdown position, which causes the model to rapidly fall forward, resulting in a failed gait transition.

Finally, Fig. 9D shows the duration when the trajectories are below the stable manifold, normalized by the gait cycle duration. This implies that when the forward step length is too large, the model falls backward due to saddle instability, resulting in a failed gait transition. As described above, there are a few exceptions where the gait transition is successful while the forward step length is above the specific range. This is because, unlike the example in Fig. 8E, the trailing limb left the ground soon after the large step of the trigger limb and then the model fell backward rapidly, fulfilling the first and second criteria.

## Discussion

In this study, we investigated the mechanism of dynamic gait transition from quadrupedal to bipedal locomotion in Japanese macaques through forward dynamics simulation using a neuromusculoskeletal model that integrates a detailed musculoskeletal framework with a motor control model based on neurophysiological findings. Through dynamical systems analysis using an inverted pendulum model, we found that an inverted pendulum motion with appropriate control of the forward step length of the trigger limb is crucial in the gait transition.

Quadrupedal walking is relatively stable because the body is supported by the four limbs, but bipedal walking is very unstable because the body is supported only by the hindlimbs, and it is easy to fall over. This instability is due to the fact that the support base is small and the motion is governed by the saddle dynamics, which has an unstable equilibrium point like an inverted pendulum. This dynamical system has the property of diverging quickly and can easily generate large movements even without control input. Japanese macaques that have been trained to walk bipedally successfully use this dynamic characteristic to achieve efficient bipedal walking [16, 29, 31]. Specifically, the rotational motion produced by the inverted pendulum system supported by one hindlimb is used as a translational motion, and the other hindlimb touches the ground before falling over to start a new inverted pendulum motion. By repeating this, bipedal walking continues. Our results suggest that Japanese monkeys also use the inverted pendulum dynamics in the transition from quadrupedal to bipedal locomotion. Specifically, from the moment the trunk is raised and bipedal walking begins, the inverted pendulum dynamics is employed and maintained, enabling a continuous and stable shift from quadrupedal to bipedal locomotion. In particular, an appropriate control of the touchdown position through a large step of the trigger limb plays a key role in the success of this process. This mechanism is reminiscent of the foot-placement control strategy proposed by Raibert [36] for spring-loaded inverted pendulum (SLIP) models and hopping robots, in which the touchdown location is regulated to control locomotor stability and forward speed. The ability to exploit inverted pendulum dynamics not only during bipedal walking but also during gait transitions could have played a role in the evolution of bipedalism.

In addition to the adequate control of touchdown position to produce a proper inverted pendulum motion, adequate coordination of the limbs and trunk is important in the gait transition due to the complexity of the musculoskeletal system in Japanese macaques. To meet this, we constructed a motor control model based on the two-layer CPG model [4, 38] and the muscle synergy hypothesis [7, 10, 13, 20, 47]. In addition, we parameterized the motor control model and created various movements, including failed gait transitions such as forward and backward falls. By comprehensively evaluating these various movements, we identified the generation of appropriate inverted pendulum motion through control of the forward step length of the trigger limb as a key mechanism underlying successful gait transitions. Such a systematic evaluation, particularly including failed gait transitions, is difficult to perform experimentally in animals and represents an important advantage of model-based simulation.

Humans also use the inverted pendulum mechanism for efficient bipedal walking [5]. Trained Japanese macaques use the inverted pendulum mechanism, but the CoM trajectory and ground reaction force pattern differ from those of humans, suggesting that monkey bipedal walking is not as efficient as human bipedal walking [29, 32]. Because Japanese macaques, which are naturally quadrupedal, walk bipedally with their trunk upright, their hip joints are extended by about 90 degrees compared to when they walk quadrupedally. Their inefficient bipedal walking may be due to anatomical limitations in the range of motion of the hip joints [33]. Such hip joint limitations should also affect the gait transition, and we would like to take this into account in the future.

In this study, we focused on the trigger limb as the primary controller of the gait transition (Fig. 4). However, the trailing limb and the forelimbs would also play an important role in this process. We would like to extend our model in the future to investigate their contributions. Nevertheless, it is worth noting that our results demonstrate that controlling the trigger limb during a single step is sufficient to enable the gait transition, with the underlying mechanism being rooted in the inverted pendulum dynamics. Our results also generate testable predictions. In particular, impairing the proper touchdown position of the trigger limb and the inverted pendulum motion should impede the gait transition.

Although we reproduced quadrupedal walking, bipedal walking, and the transition from quadrupedal to bipedal walking using a forward dynamic simulation of a neuromusculoskeletal model of the Japanese macaque, this does not mean that the exact movements observed in the real Japanese macaque were fully reproduced. For example, the ratio of swing phase and stance phase is different, and the double stance phase is short. The angle of the thigh is small, and the trunk during bipedal walking is not as upright as in the Japanese macaque. The forward step length of the trigger limb at the transition is short. In addition, the large activity of the TA and SO muscles at touchdown during the transition, which would increase ankle joint stiffness through co-contraction, is not reproduced in the model. There are several reasons for these discrepancies, including the replacement of complex musculoskeletal and motor control systems with simple models, and the computational problems that arise when model parameters are determined by optimization. Nevertheless, our model qualitatively captures the kinematics and muscle activation patterns during both quadrupedal and bipedal walking, as well as the changes during the gait transition. Therefore, it is reasonable to use the model to discuss the transition mechanism, particularly with respect to the appropriate touchdown position of the trigger limb and the dynamics of an inverted pendulum.

In addition to the quantitative discrepancy between simulation results and measured monkey data, our model has limitations. For example, we focused on locomotor behavior in the sagittal plane; however, real monkeys move in three-dimensional space, and locomotor behaviors in the frontal and transverse planes are also important. In particular, while Japanese macaques, like humans, walk facing forward during bipedal locomotion, they walk with their body axis at an angle to the walking direction during quadrupedal locomotion, as observed in rhesus macaques [6, 39]. This differs from quadrupedal locomotion in other mammals, such as cats [18], and is thought to prevent interference between the forelimb and the ipsilateral hindlimb [24]. However, it has been suggested that this particular posture also contributes to balance control during walking [15]. Furthermore, we developed a motor control model mainly based on the muscle synergy hypothesis [7, 10, 13, 20, 47], where command signals were generated by a linear combination of a small number of rectangular pulses (Figs. 2A and 4). Therefore, the waveform of the command signal for each muscle is very simple and limited. In addition, the waveform and burst timing were determined in a feedforward manner. To generate diverse movement patterns, feedback control based on sensory information, including spinal reflexes, plays an important role [9, 45]. Although we used a feedback control to regulate the waveform of the command signal in our model (Fig. 3), the contribution was very small. These limitations suggest directions for future improvement, including modeling locomotion in the frontal and transverse planes, joint stiffness control, and sophisticated feedback control. Furthermore, the present study focused on quadrupedal walking, bipedal walking, and the transition from quadrupedal to bipedal walking on level ground. Future studies should investigate whether the proposed mechanisms, including the role of inverted-pendulum dynamics, can also explain locomotor behaviors under different task and environmental conditions, such as transitions from bipedal to quadrupedal walking and gait adaptations on inclined surfaces. Such analyses would contribute to a more comprehensive understanding of the mechanisms underlying gait generation and gait transitions across a wide range of locomotor contexts.

## Methods

### Modeling

#### Musculoskeletal model

We developed a two-dimensional musculoskeletal model of Japanese monkeys based on our previous rat model [48]. The skeletal part of our model consists of eleven rigid links representing the trunk (including the head), forelimbs (two links), and hindlimbs (three links), as shown in Fig. 1. When the trunk is parallel or vertical to the ground, the angle is 0° or 90°, respectively. When the upper arm and forearm are in a straight line and parallel to the trunk, the shoulder and elbow angles are 180°. When the thigh, shank, and dorsum of the foot are in a straight line and parallel to the trunk, the hip, knee, and ankle angles are 180°. When the joints extend, the joint angles increase. The ground reaction force is modeled using one viscoelastic element at the tip of each arm and four elements at each sole. The physical parameters of the skeletal model are determined from measured data of Japanese monkeys [30] as shown in Table 1. We derived the equations of motion using Lagrangian equations and solved them using the fourth-order Runge-Kutta method with a time step of 0.05 ms.

**Table 1:** Physical parameters of skeletal model.

| Parameter | Trunk | Upper arm | Forearm | Thigh | Shank | Foot |
| --- | --- | --- | --- | --- | --- | --- |
| Mass [g] | 6517 | 320 | 261 | 557 | 268 | 101 |
| Length [mm] | 370 | 155 | 230 | 163 | 182 | 74 |
| MOI [$\times 10^5 \text{g} \cdot \text{mm}^2$] | 1720 | 5.96 | 13.8 | 16.1 | 7.01 | 1.21 |

MOI: Moment of inertia around CoM

For the muscle part of our model, we used six principal muscles for each forelimb: four uniarticular, namely shoulder flexor (anterior deltoid, AD), shoulder extensor (posterior deltoid, PD), elbow flexor (brachioradialis, BR), and elbow extensor (anconeus, AN), and two biarticular, namely shoulder flexor and elbow flexor (biceps brachii, BB) and shoulder extensor and elbow extensor (triceps brachii, TB), as shown in Fig. 1. We used eight principal muscles for each hindlimb: five uniarticular, namely hip flexor (iliopsoas, IP), hip extensor (gluteus maximus, GM), knee extensor (vastus lateralis, VL), ankle flexor (tibialis anterior, TA), and ankle extensor (soleus, SO), and three biarticular, namely hip flexor and knee extensor (rectus femoris, RF), hip extensor and knee flexor (biceps femoris, BF), and knee flexor and ankle extensor (gastrocnemius, GA). The moment arms of the muscles around the joints are constant, regardless of joint angles. We modeled muscle tension *F*_*m,i*_ (*m* = AD, PD, BR, AN, BB, TB, IP, GM, VL, TA, SO, RF, BF, GA, *i* = left, right) through contractile and passive elements based on the previous work [12, 48] by

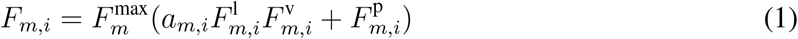

where 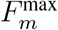 is the maximum muscle tension, *a*_*m,i*_ is the muscle activation (*a*_*m,i*_ ≥ 0), 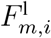 is the force-length relationship, 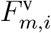 is the force-velocity relationship, and 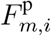 is the passive component. The muscle lengths are normalized by 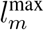, which is set so that all uniarticular muscles had a length of 85% of 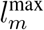 and all biarticular muscles have a length of 75% of 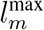 at a neutral posture with the shoulder at 70°, the elbow at 45°, the hip at 70°, the knee at 90°, and the ankle at 80°. Furthermore, 2° of joint motion corresponds to 1% of muscle length change, except for GA (1.5° at the ankle or 4.5° at the knee). The muscle contractile velocities are normalized by 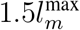. The physical parameters of the muscle model are determined from measured data of Japanese monkeys [30] as shown in Table 2.

**Table 2:** Physical parameters of muscle model.

| Parameter | AD | PD | BR | AN | BB | TB |  |  |  |
| --- | --- | --- | --- | --- | --- | --- | --- | --- | --- |
| $F_m^{\max}$ [N] | 116 | 106 | 306 | 771 | 444 | 845 | | | |
| MA [mm] | 10 | 10 | 15 | 20 | 10 (s) | 25 (s) |  |  |  |
|  |  |  |  |  | 15 (e) | 20 (e) |  |  |  |
| Parameter | IP | GM | VL | TA | SO | RF | BF | GA |  |
| $F_m^{\max}$ [N] | 644 | 739 | 2513 | 391 | 824 | 718 | 803 | 718 | |
| MA [mm] | 20 | 40 | 25 | 15 | 20 | 20 (h) | 40 (h) | 10 (k) |  |
|  |  |  |  |  |  | 25 (k) | 40 (k) | 20 (a) |  |

MA: moment arm of muscle around joint, s: shoulder, e: elbow, h: hip, k: knee, and a: ankle

The muscle activation *a*_*m,i*_ is determined through the low-pass filter given by

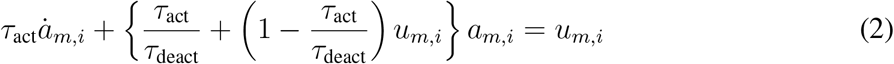

where *τ*_act_ (= 28 ms) and *τ*_deact_ (= 44 ms) are activation and deactivation time constants, respectively, and *u*_*m,i*_ is the motor command determined in the motor control model.

#### Motor control model

We developed a motor control model based on the hypothetical motor program at the spinal cord level (movement generator) and the regulation of locomotion movement at the brainstem and cerebellum levels (movement regulator) in the same way as that in our previous work [2, 3, 48]. The movement generator produces motor commands in a feedforward fashion to create limb movements based on the muscle synergy hypothesis. The movement regulator creates motor commands to regulate locomotor behavior in a feedback fashion based on sensory information.

For the movement generator, we focused on the hypothetical two-layer CPG model composed of a rhythm generator (RG) network, which produces rhythm and phase information for motor commands, and a pattern formation (PF) network, which produces spatiotemporal patterns of motor commands [4, 38]. For the RG model, we used four phase oscillators, each of which produces a basic rhythm and phase information for the corresponding limb. The phase is given by 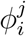 (*i* = left, right, *j* = fore, hind; 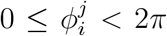) and follows 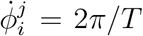, where *T* is the gait cycle duration (0.76 s for quadrupedal and 0.6 s for bipedal walking based on the monkey data measured in our previous work [15]). The oscillators remain in antiphase between the left and right sides and between the fore and hind sides. For the PF model, we determined the motor commands to generate limb movements using the corresponding oscillator phase based on the muscle synergy hypothesis, which suggests that the linear combination of only a small number of basic signals produces a large portion of motor commands in animal locomotion [10, 20, 23, 25, 37]. Specifically, we used four rectangular pulses 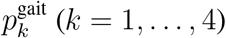 for quadrupedal and bipedal walking, which are given by

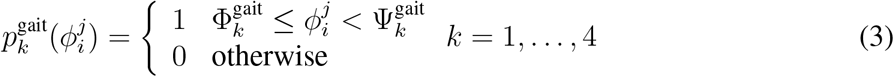

where 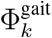 and 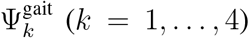 are the onset and end phases of the pulse 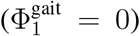, respectively, and used the same value between the forelimbs and hindlimbs. Specifically, 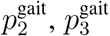, and 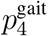 contribute to early flexion, late flexion, early extension, and late extension, respectively, where 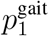 uses the PD, BR, and, BB muscles for the forelimbs and IP, TA, and BF muscles for the hindlimbs, 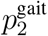 uses the AD, BR, and, BB muscles for the forelimbs and IP, VL, and TA muscles for the hindlimbs, 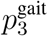 uses the AD, BR, AN, BB, and TB muscles for the forelimbs and GM, VL, SO, RF, BF, and GA muscles for the hindlimbs, and 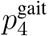 uses the PD, BR, AN, BB, and TB muscles for the forelimbs and GM, VL, SO, RF, BF, and GA muscles for the hindlimbs, as shown in Fig. 2. The motor command 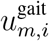 of the movement generator for quadrupedal and bipedal walking is given by

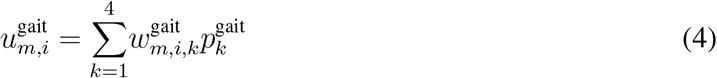

where 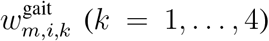 is the weighting coefficient. This movement generator has 41 parameters to be determined (three for 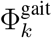, four for 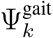, and 34 for 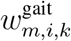).

For the movement regulator, we focused on the regulation of locomotor behavior based on sensory information at the levels of the brainstem and cerebellum. Specifically, we regulated the forward speed, hip height, and trunk posture using simple feedback controls of the muscles of standing hindlimbs, as shown in Fig. 3. For the forward speed, we used the IP, GM, TA, and SO muscles to maintain the speed. The motor command 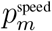 (*m* = IP, GM, TA, SO) is given by

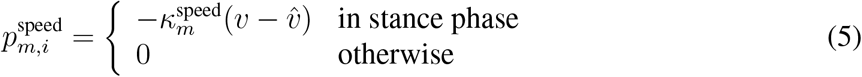

where *v* is the forward speed, 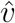 is its desired value, and 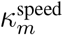 is the gain parameter (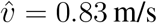 based on the monkey data measured in our previous work [15]). For the hip height, we used the VL, TA, and SO muscles to maintain the hip height. The motor command 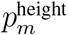 (*m* = VL, TA, SO) is given by

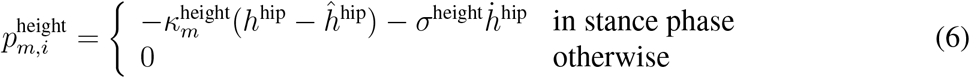

where *h*^hip^ and 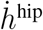 are the hip height and its rate, respectively, *ĥ*^hip^ is the reference height, and 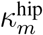 and 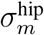 are the gain parameters. For the trunk posture, we used the IP and GM muscles to maintain the trunk angle only during bipedal walking. The motor command 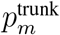 (*m* = IP, GM) is given by

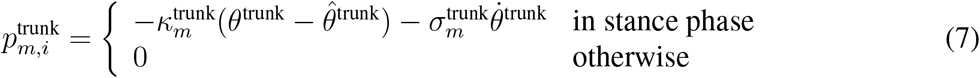

where *θ*^trunk^ and 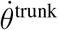 are the trunk angle and its rate, respectively, 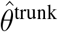 is the reference angle, and 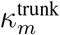and 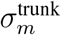 are the gain parameters. The motor command 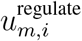 of the movement regulator is given by

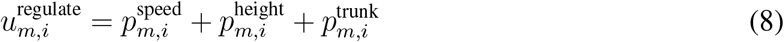

This movement regulator has 11 parameters for quadrupedal walking and 16 parameters for bipedal walking to be determined (one for *ĥ*^hip^ and 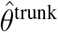, four for 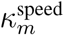, three for 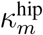 and 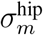, and two for 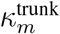 and 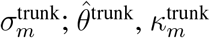, and 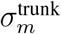 are only for bipedal walking).

To change from quadrupedal to bipedal walking, we added two rectangular pulses 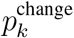 (*k* = 1, 2) for the right hindlimb (trigger limb) only once to the movement generator, which are given by

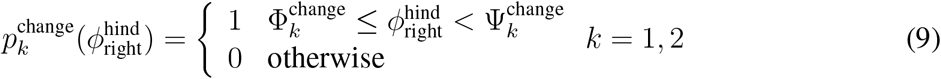

where 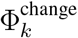 and 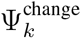 (*k* = 1, 2) are the onset and end phases of the pulse, respectively. Specifically, 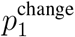 and 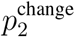 contribute to stepping the right hindlimb (trigger limb) forward and raising the trunk, respectively, where 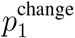 uses the IP and VL muscles and 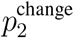 uses the GM and BF muscles, as shown in Fig. 4. The model first walked quadrupedally using the parameters for quadrupedal walking. After 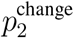 was added, we changed the values of the gait cycle, 41 parameters of the movement generator (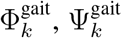, and 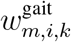), and 11 parameters of the movement regulator (*ĥ*^hip^, 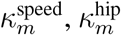, and 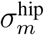) from quadrupedal to bipedal walking in both left and right sides, and used five parameters of the movement regulator for bipedal walking (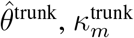, and 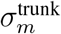). The motor command 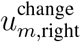 for this gait transition is given by

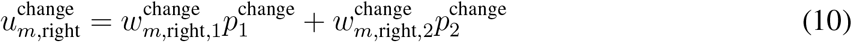

where 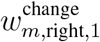 and 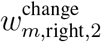 are the weighting coefficients. This additional movement generator for gait transition has eight parameters to be determined (two for 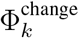, two for 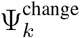, and four for 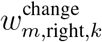).

The motor command *u*_*m,i*_ is the summation of the three components from the movement generator and regulator as follows:

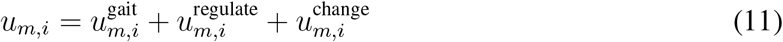

#### Parameter determination

We determined the parameters of the motor control model through forward dynamic simulations of our neuromusculoskeletal model using the PSO method [21]. Specifically, we performed simulations of bipedal walking, quadrupedal walking, and gait transition separately and determined the model parameters for each task. The initial values of the parameters to be optimized were determined based on the corresponding measured monkey data. The number of particles in the PSO was set to ten times the number of parameters to be optimized, and the optimization was performed for 50 iterations.

First, we performed the simulation of quadrupedal walking for 5.3 s (seven gait cycles) and determined the 41 parameters of the gait generator (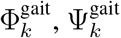, and 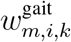) and 11 parameters of the movement regulator (*ĥ*^hip^, 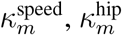, and 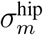) to minimize the error between the simulated and measured angles of the trunk, forelimbs, and hindlimbs during the last gait cycle. Second, we performed the simulation of bipedal walking for 4.2 s (seven gait cycles) and determined the 41 parameters of the gait generator (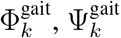, and 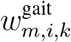) and 16 parameters of the movement regulator (*ĥ*^hip^, 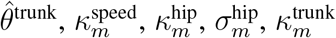, and 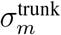) to minimize the error between the simulated angles and measured angles of the trunk and hindlimbs during the last gait cycle, where we neglected the forelimbs because the monkey forelimbs move to catch food as a reward during trials. Lastly, we performed the simulation of gait transition for 4.8 s, where 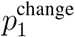 was added after three gait cycles of quadrupedal walking, and determined the eight parameters for the gait generator (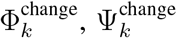, and 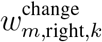) to minimize the error between the simulated and measured angles of the trunk and hindlimbs during one gait cycle after 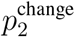 was added. That is, we evaluated the error only during bipedal walking after the gait transition. The determined parameters are shown in Table S1 for the gait generator for quadrupedal and bipedal walking, Table S2 for the gait regulator, and Table S3 for the gait generator for gait transition in Supplementary Information S1.

#### Investigation of the contribution of trigger limb using control parameters

To investigate the contribution of the trigger limb to the gait transition, we performed the simulation of gait transition using the eight parameters for the gait generator of the gait transition (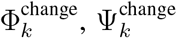, and 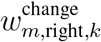). Specifically, for the first input 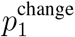, we used the same value for the onset phase 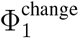 and the end phase 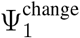 as in Table S3, and systematically varied 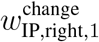 and 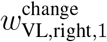 between 0 and 1. On the other hand, for the second input 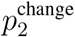, we determined 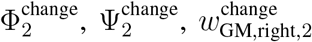, and 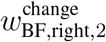 by the same optimization as above. To evaluate the contribution of the trigger limb, we considered the gait transition successful if the CoM height averaged over one gait cycle after adding 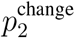 reached 0.4 m based on that during bipedal walking (first criterion), and if the trigger limb lifted off the ground within one gait cycle (0.6 s) after touchdown to continue walking (second criterion). These criteria were normalized by the hindlimb length (0.42 m) and the gait cycle duration (0.6 s), respectively.

#### Analysis using inverted pendulum model

When the trunk is raised during the gait transition, the forelimbs no longer support the body and bipedal walking with alternating left-right hindlimb stepping starts. To investigate the mechanism of gait transition, we used an inverted pendulum model with angle *θ* (Fig. 8A), which connects from the touchdown position of the trigger limb to the CoM. Specifically, we calculated the CoM position and the touchdown position of the trigger limb from the neuromusculoskeletal model simulation and the measured monkey data to determine *θ* and 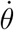.

To investigate whether the neuromusculoskeletal model simulation results and the measured monkey data show inverted pendulum motion, we compared their trajectories of (*θ*, 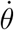)with those obtained from the equation of motion of the inverted pendulum model 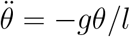, where *l* is the pendulum length and *g* is the acceleration of gravity. Specifically, we numerically solved the equations of motion during the single stance phase of the hindlimbs so that the trajectories pass through the state (*θ*, 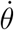) obtained at the midpoint of the single stance phase of the neuromusculoskeletal model simulation and measured monkey data. We then calculated the root mean square error between the trajectory of the neuromusculoskeletal model simulation or the measured monkey data and that of the inverted pendulum model simulation. These trajectories were nondimensionalized using 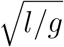, where *l* was defined as the average length during the single stance phase. In addition, these trajectories were calculated based on hindlimb support, regardless of forelimb support.

### Monkey data measurement

To verify the simulation results, we compared them with kinematic and EMG data measured from an adult Japanese monkey (*Macaca fuscata*, male, age 7 years, weight 10.4 kg) trained for 3 years to walk quadrupedally or bipedally, and to walk freely without any restraint or support on a treadmill (ORK-2000; Ohtake-Root Kogyo, Ichinoseki, Japan) at a belt speed of 0.83–1.0 m/s. The experiments were reviewed and approved by the Animal Care and Use Committee of Kindai University and were performed in accordance with the Guidelines for Proper Conduct of Animal Experiments of the Science Council of Japan. Because the details of the experimental setup and design are explained in our previous work [15], we briefly explain them.

The monkey walked quadrupedally or bipedally and changed the gait several times on the treadmill at 1.0 m/s of the belt speed during a trial of 60 s. They were filmed with high-speed video cameras (HAS-220; DITECT, Tokyo, Japan) at 200 frames/s and with markers over the following 20 landmarks: tragus, external occipital protuberance, and base of the tail for the body; and left and right acromion (shoulder), lateral epicondyle of the humerus (elbow), styloid process of the ulna (wrist), fifth metacarpophalangeal joint, greater trochanter of the femur (hip), lateral epicondyle of the femur (knee), lateral malleolus of the fibula (ankle), and fifth metatarsophalangeal joint for the limbs. To estimate the CoM during walking, we used the anatomical mass distribution estimated in [30], assuming that the mass of the forelimbs is concentrated on the trunk during bipedal walking. EMG activity was recorded using implanted electrodes and a main amplifier (MEG-6116; Nihon Kohden) at a sampling rate of 10 kHz from the right forelimb and right hindlimb muscles and from the trunk muscles in both sides of the body. Specifically, 13 muscles were recorded; the BB, TB, flexor carpi ulnaris, and extensor carpi radialis muscles from the forelimb, the erector spinae and contralateral erector spinae muscles from the trunk, and the GM, TA, SO, RF, BF, GA, and flexor hallucis longus muscles from the hindlimb. EMG data from two forelimb muscles (BB and TB muscles) and six hindlimb muscles (GM, TA, SO, RF, BF, and GA muscles) were used for the comparison with the simulation results. These EMG data were full-wave rectified, smoothed, and normalized by the maximum value during quadrupedal, bipedal, and transition in each muscle.

We measured the activity of the forelimb and hindlimb muscles only on the right side of the body. Since the roles of the right and left muscles were considered to be almost identical during quadrupedal and bipedal walking, we compared the measured data with the simulation results. However, the roles of the right and left muscles differed during the transition from quadrupedal to bipedal walking, because a hindlimb (trigger limb) took a large step and the trunk was then raised. This means that muscle activity is required on both sides of the body for comparison. However, in some cases, it was the right hindlimb that acted as the trigger limb, while in others it was the left. Therefore, when the right hindlimb acted as the trigger limb, we used the measured data as the trigger limb. In contrast, when the left hindlimb acted as the trigger limb, we used the measured data as the trailing limb. This enabled us to compare the measured data with the simulation results during the transition.

## Acknowledgments

This study was supported in part by JSPS KAKENHI Grant Numbers JP24H00297, JP20H00229, and JP19KK0377, and JST FOREST Program Grant Number JPMJFR2021.

## Data availability

The data will be made available upon acceptance of the manuscript.

## Author contributions

SA developed the study design. KoN and YT carried out the computer simulations of the monkey model. TS, YH, KM, and KaN performed the experiments on the Japanese monkey. KoN and HO analyzed the data in consultation with MA, YA, RI, and SA. KoN and SA wrote the manuscript and all the authors reviewed and approved it.

