## Supplementary Information for "Gait transition mechanism from quadrupedal to bipedal locomotion in the Japanese macaque based on inverted pendulum"

### S1. Optimization results

Tables S1, S2, and S3 show the parameters determined through optimization for the movement generator for quadrupedal and bipedal walking, the movement regulator, and the movement generator for gait transition, respectively.

Table S1: Determined parameters of movement generator for quadrupedal and bipedal walking.

| | Quadrupedal | | | | Bipedal | | | |
|---|---|---|---|---|---|---|---|---|
| | 1 | 2 | 3 | 4 | 1 | 2 | 3 | 4 |
| $\Phi_k^{\text{gait}}$ [rad] | 0 | 0.71 | 2.42 | 3.64 | 0 | 0.69 | 1.87 | 3.09 |
| $\Psi_k^{\text{gait}}$ [rad] | 0.63 | 1.62 | 3.48 | 5.26 | 0.66 | 1.64 | 2.92 | 4.68 |
| | $w_{m,i,1}^{\text{gait}}$ | $w_{m,i,2}^{\text{gait}}$ | $w_{m,i,3}^{\text{gait}}$ | $w_{m,i,4}^{\text{gait}}$ | $w_{m,i,1}^{\text{gait}}$ | $w_{m,i,2}^{\text{gait}}$ | $w_{m,i,3}^{\text{gait}}$ | $w_{m,i,4}^{\text{gait}}$ |
| AD | - | 0.61 | 0.21 | - | - | 0.00 | 0.00 | - |
| PD | 0.16 | - | - | 0.22 | 0.03 | - | - | 0.03 |
| BR | 0.05 | 0.05 | 0.05 | 0.05 | 0.01 | 0.02 | 0.01 | 0.05 |
| AN | - | - | 0.22 | 0.22 | - | - | 0.00 | 0.01 |
| BB | 0.05 | 0.05 | 0.05 | 0.05 | 0.03 | 0.03 | 0.03 | 0.03 |
| TB | - | - | 0.16 | 0.16 | - | - | 0.03 | 0.01 |
| IP | 0.53 | 0.32 | - | - | 0.42 | 0.44 | - | - |
| GM | - | - | 0.09 | 0.14 | - | - | 0.22 | 0.24 |
| VL | - | 0.28 | 0.08 | 0.17 | - | 0.08 | 0.15 | 0.21 |
| TA | 0.12 | 0.14 | - | - | 0.11 | 0.50 | - | - |
| SO | - | - | 0.15 | 0.20 | - | - | 0.69 | 0.61 |
| RF | - | - | 0.05 | 0.24 | - | - | 0.07 | 0.24 |
| BF | 0.10 | - | 0.16 | 0.12 | 0.05 | - | 0.18 | 0.89 |
| GA | - | - | 0.18 | 0.13 | - | - | 0.44 | 0.93 |

### S2. Sensitivity analysis

We performed a sensitivity analysis of the eight parameters determined through optimization for the gait transition ($\Phi_1^{\text{change}}$, $\Psi_1^{\text{change}}$, $\Phi_2^{\text{change}}$, $\Psi_2^{\text{change}}$, $w_{\text{IP,right},1}^{\text{change}}$, $w_{\text{VL,right},1}^{\text{change}}$, $w_{\text{GM,right},2}^{\text{change}}$, and $w_{\text{BF,right},2}^{\text{change}}$). Specifically, each parameter was independently varied from its optimized value by $-p$, 0, or

Table S2: Determined parameters of movement regulator.

| | Quadrupedal | | Bipedal | |
|---|---|---|---|---|
| | $\kappa_m^{\text{speed}}$ | | $\kappa_m^{\text{speed}}$ | |
| IP | -0.28 | | -0.10 | |
| GM | 0.28 | | 0.08 | |
| TA | -0.20 | | -0.10 | |
| SO | 0.20 | | 0.08 | |
| $\hat{h}^{\text{hip}}$ [m] | 0.36 | | 0.30 | |
| | $\kappa_m^{\text{hip}}$ | $\sigma_m^{\text{hip}}$ | $\kappa_m^{\text{hip}}$ | $\sigma_m^{\text{hip}}$ |
| VL | 1.00 | 0.0040 | 1.19 | 0.0002 |
| TA | -1.01 | -0.0040 | -1.50 | -0.0002 |
| SO | 1.00 | 0.0040 | 0.81 | 0.0002 |
| $\hat{\theta}^{\text{trunk}}$ [°] | - | | 0.8 | |
| | $\kappa_m^{\text{trunk}}$ | $\sigma_m^{\text{trunk}}$ | $\kappa_m^{\text{trunk}}$ | $\sigma_m^{\text{trunk}}$ |
| IP | - | - | -0.54 | -0.09 |
| GM | - | - | 0.71 | 0.58 |

$+p\%$, resulting in $3^8$ ($= 6561$) combinations. Gait-transition simulations were performed for all combinations, and the success or failure of the gait transition was evaluated based on the two criteria used in Fig. 7: the CoM height averaged over one gait cycle after adding two pulses and the time from touchdown to liftoff of the trigger limb following a large step. This analysis was performed for $p = 5$, 10, 15, and 20.

Figure S1A shows the success rates calculated using all trials and those calculated separately according to the number of parameters varied from their optimized values. The success rate decreased as the magnitude of the parameter variation increased and as the number of varied parameters increased. However, the success rate did not decrease abruptly for small parameter variations and remained relatively high over a range of parameter variations, indicating that successful gait transitions were maintained despite perturbations of the optimized parameters. Figure S1B shows the success rate associated with positive and negative variations of each individual parameter. For each parameter and each variation ($-p$ or $+p\%$), the success rate was

Table S3: Determined parameters of movement generator for gait transition.

| | 1 | 2 |
|---|---|---|
| $\Phi_k^{\text{change}}$ [rad] | 1.48 | 3.30 |
| $\Psi_k^{\text{change}}$ [rad] | 2.56 | 6.18 |
| | $w_{m,\text{right},1}^{\text{change}}$ | $w_{m,\text{right},2}^{\text{change}}$ |
| IP | 0.41 | - |
| VL | 0.29 | - |
| GM | - | 1.00 |
| BF | - | 0.85 |

calculated from the $3^7$ $(= 2187)$ trials in which that parameter was fixed at the specified variation while the other seven parameters independently took values of $-p$, 0, or $+p\%$. The success rate was particularly affected by variations in $\Psi_1^{\text{change}}$, $\Psi_2^{\text{change}}$, and $w_{\text{IP},\text{right},1}^{\text{change}}$. Nevertheless, no single parameter variation caused an abrupt decrease in the success rate, suggesting that the gait-transition simulation was not critically dependent on a specific optimized parameter.

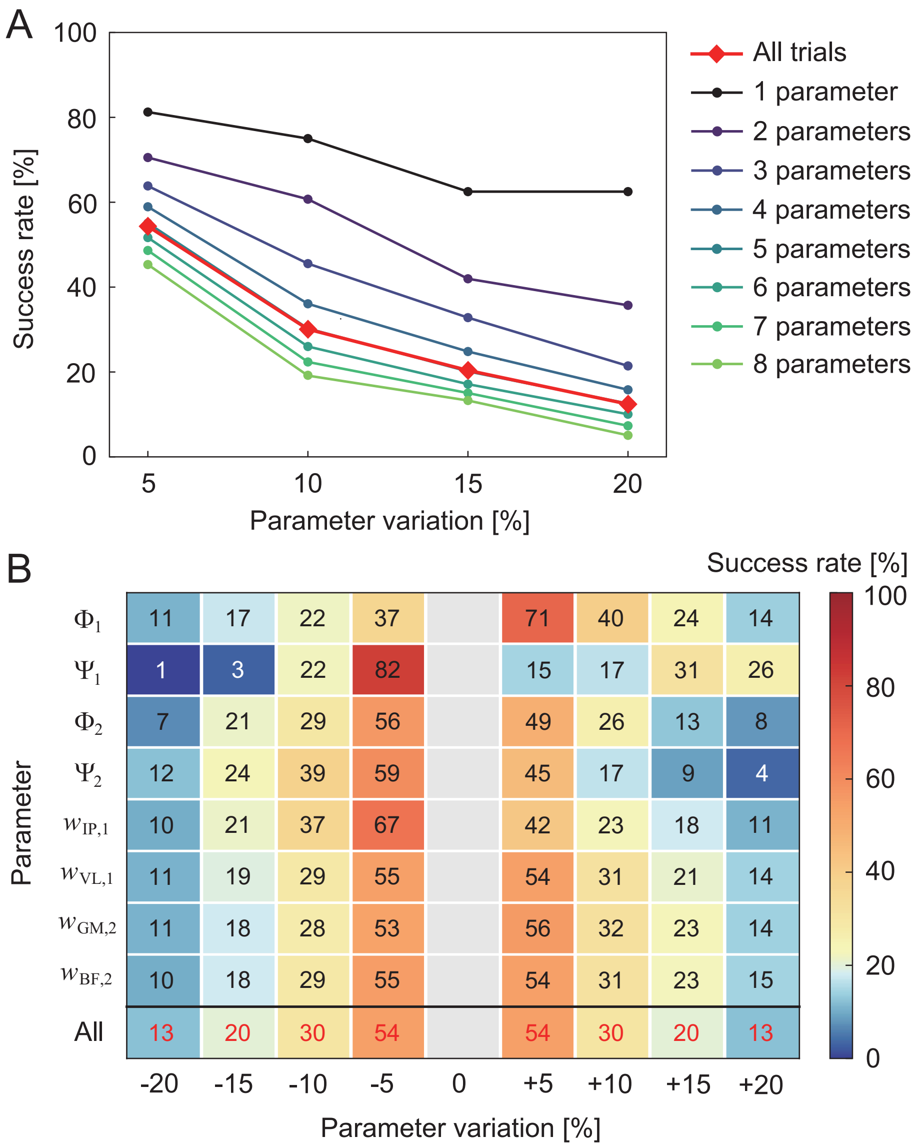


Figure S1: Sensitivity analysis of the optimized parameters for gait transition. A. Success rates calculated using all trials and separately according to the number of parameters varied from their optimized values. B. Success rates associated with positive and negative variations of each individual parameter. The 0% variation is not shown because it does not represent a perturbation of the individual parameter and therefore does not provide a direct measure of its sensitivity. The bottom row shows the success rates calculated using all trials for each magnitude of parameter variation.
